# SIDT2 RNA transporter promotes lung and gastrointestinal tumor development

**DOI:** 10.1101/724237

**Authors:** Tan A Nguyen, Kathryn T Bieging-Rolett, Tracy L Putoczki, Ian P Wicks, Laura D Attardi, Ken C Pang

## Abstract

RNautophagy is a newly-described type of selective autophagy whereby cellular RNAs are transported into lysosomes for degradation. This process involves the transmembrane protein SIDT2, which transports double-stranded RNA (dsRNA) across the endolysosomal membrane. We previously demonstrated that SIDT2 is a transcriptional target of *p53*, but its role in tumorigenesis – if any - is unclear. Unexpectedly, we show here that *Sidt2^−/−^* mice with concurrent oncogenic *Kras*^G12D^ activation develop significantly fewer tumors than littermate controls in a mouse model of lung adenocarcinoma (LUAD). Consistent with this observation, loss of SIDT2 also leads to enhanced survival and delayed tumor development in an *Apc*^*min*/+^ mouse model of intestinal cancer. Within the intestine, *Apc*^*min*/+^;*Sidt2*^−/−^ mice display accumulation of dsRNA in association with increased phosphorylation of eIF2α and JNK as well as elevated rates of apoptosis. Taken together, our data demonstrate a role for SIDT2 - and by extension RNautophagy - in promoting tumor development.

## INTRODUCTION

The *C. elegans* double-stranded RNA transporter (dsRNA) SID-1 is conserved throughout much of animal evolution, and mammals possess two paralogs, SIDT1 and SIDT2 ^1,2^. SIDT2 is broadly expressed in mammalian tissue and localizes within late endosomes and lysosomes ^3,4^. Human and mouse *Sidt2* homologues show 95% sequence identity across the entire protein (832 amino acids) and 100% identity at the C-terminal 100 amino acids ^4^. Such a high degree of conservation implies a strongly-selected function, and studies have recently emerged that shed light on the role of SIDT2 in mammals.

On the one hand, SIDT2 appears to have retained RNA transporter activity. This was initially suggested by the observation that the ectodomain of SIDT2 binds long dsRNA *in vitro*, similar to *C. elegans* SID-1 ^5^. Consistent with this finding, we subsequently discovered that SIDT2 transports viral dsRNA and that this transport is important for anti-viral immunity ^4^. More specifically, we found that SIDT2 promotes the trafficking of internalized dsRNA across the endo-lysosomal membrane and into the cytoplasm, where it is recognized by RNA sensors, which in turn promote anti-viral, type I interferon (IFN) signaling. Loss of SIDT2 thus impairs IFN production and survival after viral infection is significantly reduced ^4^. In parallel, SIDT2 has also recently been reported to traffic RNAs *into* the lysosome for degradation in a novel process described as “RNautophagy” ^6^. These experiments - performed using cell-free biochemical assays - suggested that SIDT2 promotes destruction of endogenous RNAs by transporting them from the cytosol into lysosomes. Such transport would thus be in the opposite direction to that described for viral RNAs, but is potentially consistent with previous observations that RNA transport by *C. elegans* SID-1 is bidirectional and dependent on RNA concentration ^1^.

On the other hand, some studies have observed physiological effects of SIDT2 where the relationship to RNA transport – if any - is unclear. For example, mice lacking SIDT2 demonstrate impaired glucose tolerance, decreased serum insulin levels, and defective insulin secretion ^7–9^. Two recent studies also demonstrated that *Sidt2*^−/−^ mice develop non-alcoholic fatty liver disease (NAFLD) ^10,11^, with one suggesting that this is due to induction of endoplasmic reticulum stress ^10^ and the other proposing that it is the result of defective autophagy ^11^. Finally, work from our group has also demonstrated a potential role for SIDT2 in tumorigenesis ^12^. Specifically, we found that SIDT2 is a transcriptional target of the tumor suppressor p53, that SIDT2 overexpression in *HrasV12;p53* null mouse embryo fibroblasts (MEFs) impairs cell proliferation, and that shRNA-mediated knockdown of *Sidt2* in a fibrosarcoma model leads to increased tumor growth following transplantation into immunocompromised *Scid* mice ^12^. Together with the observation that *SIDT2* is transcriptionally downregulated in patient tumors compared to healthy tissue ^13^, these findings thus support a possible tumor suppressive role for SIDT2.

In the current study, we further investigated the role of SIDT2 in tumor development. Unexpectedly, we found that mice lacking SIDT2 display reduced tumor burden and increased survival in both lung adenocarcinoma (LUAD) and intestinal cancer models. Moreover, consistent with its role in dsRNA transport, loss of SIDT2 leads to accumulation of dsRNA, resulting in increased phosphorylation of eIF2α and elevated rates of apoptosis. Our findings therefore suggest that SIDT2 - and by extension RNautophagy - play a role in promoting tumor development.

## RESULTS

### Loss of SIDT2 inhibits lung adenocarcinoma development

Given the finding that *Sidt2* is a p53 target gene, we sought to investigate its role in tumor suppression *in vivo*. Lung cancer is the leading cause of cancer deaths worldwide, and loss or mutation of *p53* is common in this tumor type. Therefore, we examined the role of Sidt2 in LUAD tumorigenesis by employing an autochthonous mouse model in which mice conditionally express oncogenic *Kras*^*G12D*^ under the control of a lox-STOP-lox element (*Kras*^*LSL-G12D*^). Intratracheal inoculation of adenoviral Cre recombinase excises the STOP cassette, resulting in expression of Kras^G12D^ specifically in lung cells. Kras^G12D^ expression drives development of non-small cell lung tumors, and loss of p53 promotes tumor progression in this model ^14^. To assess the role of SIDT2 in tumorigenesis in this LUAD model, we crossed *Sidt2*^−/−^ mice previously generated in our lab ^4^ with *Kras*^*LSL-G12D*/+^ mice, and subsequently assessed lung tumor burden in *Kras*^*LSL-G12D*/+^;*Sidt2*^+/+^ and *Kras*^*LSL-G12D*/+^;*Sidt2*^−/−^ mice eighteen weeks after intratracheal adenoviral inoculation. In contrast to our previous report suggesting that SIDT2 has a tumor suppressive role in fibrosarcoma, light microscopic analysis of H&E-stained lung sections showed that *Sidt2*^−/−^ animals have reduced tumor burden (Figure 1A). This was confirmed with subsequent quantification, which showed that mice deficient in SIDT2 developed significantly fewer tumors (Figure 1B) and had a significant reduction in overall tumor burden (Figure 1C). Next, we wanted to investigate whether the loss of SIDT2 leads to an impairment of cellular proliferation. To do so, we compared expression of Ki67, a cellular marker of proliferation, using immunohistochemical staining (Figure 1D). Consistent with an impairment in cellular proliferation, tumors from *Kras*^*LSL-G12D*/+^;*Sidt2*^−/−^ mice had significantly less Ki67-positive cells compared to controls (Figure 1E). Together, these results thus suggest that SIDT2 facilitates tumor development in the *Kras*^*G12D*^ LUAD model.

**Figure 1.**
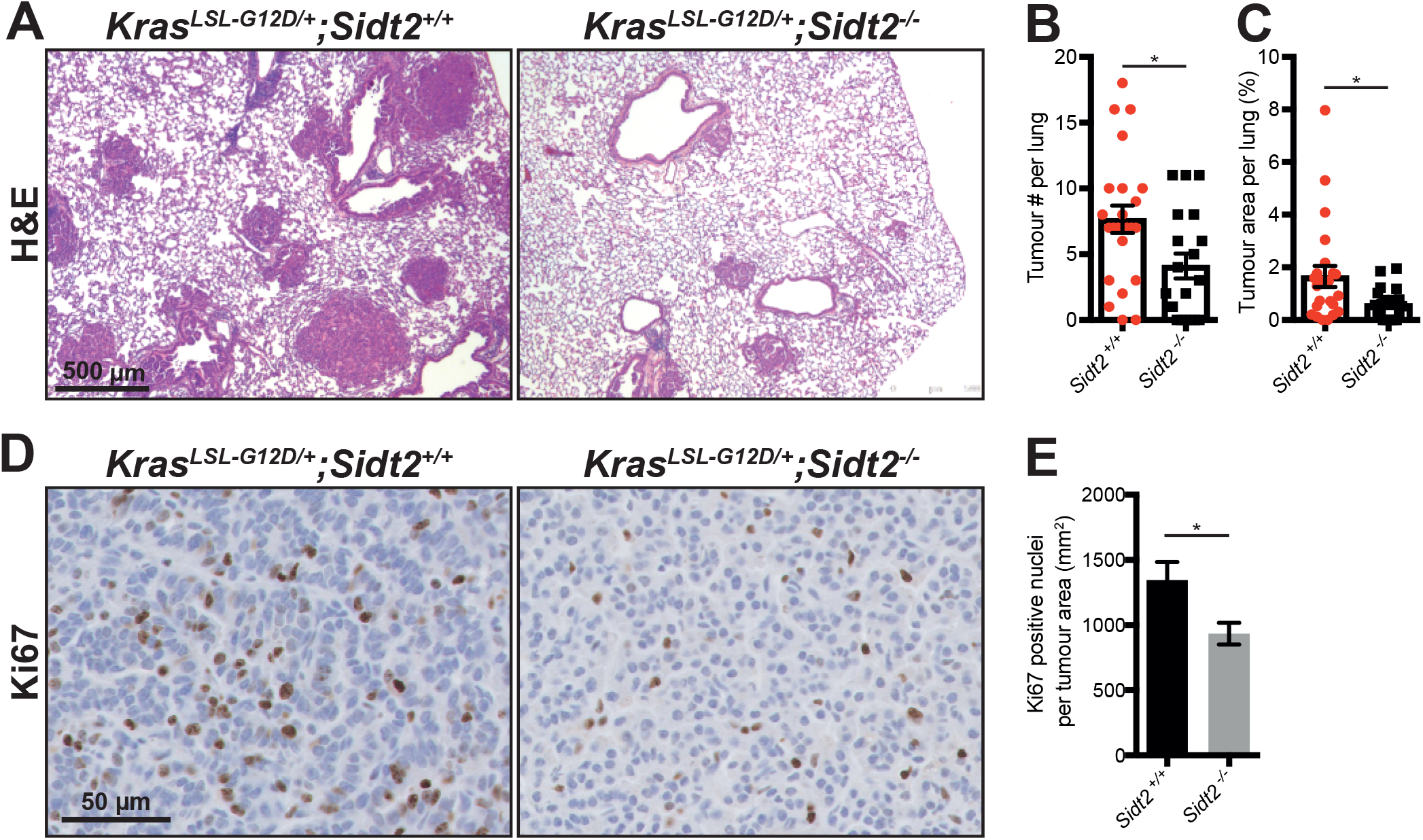
*Kras*^*LSL-G12D*/+^;*Sidt2*^−/−^ mice have increased tumor burden compared to controls. **(A)** Representative images of H&E stained lung sections from *Kras*^*LSL-G12D*/+^;*Sidt2*^+/+^ and *Kras*^*LSL-G12D*/+^;*Sidt2*^−/−^ mice eighteen weeks following inoculation with 4×10^6^ PFU Adenovirus containing Cre recombinase by intratracheal intubation. **(B)** Average tumor number and **(C)** tumor burden as a percentage of total lung area was assessed per lung section (n = 19−23 mice per genotype). **(D)** Representative images of Ki67 stained lung sections *Kras*^*LSL-G12D*/+^;*Sidt2*^+/+^ and *Kras*^*LSL-G12D*/+^;*Sidt2*^−/−^ mice. **(E)** Quantification of average number of Ki67 positive cells per mm^2^ tumor tissue. Analysis was performed on >25 tumors per mouse (n = mice per genotype). Error bars represent ± SEM. * indicates *P* < 0.05, *** indicates *P* < 0.001.

### Loss of SIDT2 inhibits growth of *Apc*^*min*^ intestinal tumors

The difference in the role for SIDT2 in tumorigenesis in the mouse fibrosarcoma and LUAD models prompted us to test another mouse tumor model to examine for context-dependency of Sidt2 in cancer. To this end, we chose the well characterized *Apc*^*min*^ mouse model of intestinal cancer. These mice harbor a dominant mutation in the oncogenic *Apc* gene which leads to spontaneous development of adenomatous polyps, primarily in the distal small intestine (DSI) ^15,16^.

We subsequently generated *Apc*^*min*/+^ mice lacking *Sidt2* (Figure S1) and monitored the animals over time, sacrificing them after the onset of anemia and/or signs of ill health associated with death in this model (e.g. hunching of the back, weight loss). *Apc*^*min*/+^;*Sidt2*^−/−^ mice (median survival: 131 days) survived significantly longer than both *Apc*^*min*/+^;*Sidt2*^+/+^ and *Apc*^*min*/+^;*Sidt2*^+/−^ mice (median survival: 93 and 99 days respectively), suggesting that loss of SIDT2 impairs intestinal tumor development (Figure 2A). To investigate further, *Apc*^*min*/+^;*Sidt2*^−/−^ mice were sacrificed at day 100 and appeared to have a lower tumor burden within the DSI compared to *Apc*^*min*/+^;*Sidt2*^+/+^ mice (Figure 2B). To properly assess this, tumor number (Figure 2C-E) and tumor area (Figure 2G-I) were calculated in the distal (DSI), medial (MSI) and proximal (PSI) small intestine of 100-day old *Apc*^*min*/+^;*Sidt2*^+/+^ and *Apc*^*min*/+^;*Sidt2*^−/−^ mice. While there was no difference in tumor number in the colon (Figure S2A) or PSI, *Apc*^*min*/+^;*Sidt2*^−/−^ mice had significantly fewer tumors than *Apc*^*min*/+^;*Sidt2*^+/+^ mice in the MSI and DSI. Moreover, *Apc*^*min*/+^;*Sidt2*^−/−^ mice also had smaller tumors than *Apc*^*min*/+^;*Sidt2*^+/+^ mice in the DSI, MSI and PSI, but again showed no difference in the colon, where tumors are uncommon (Figure S2B). Together, these results suggest that SIDT2 also facilitates tumor development in the *Apc*^*min*^ mouse model of intestinal cancer.

**Figure 2.**
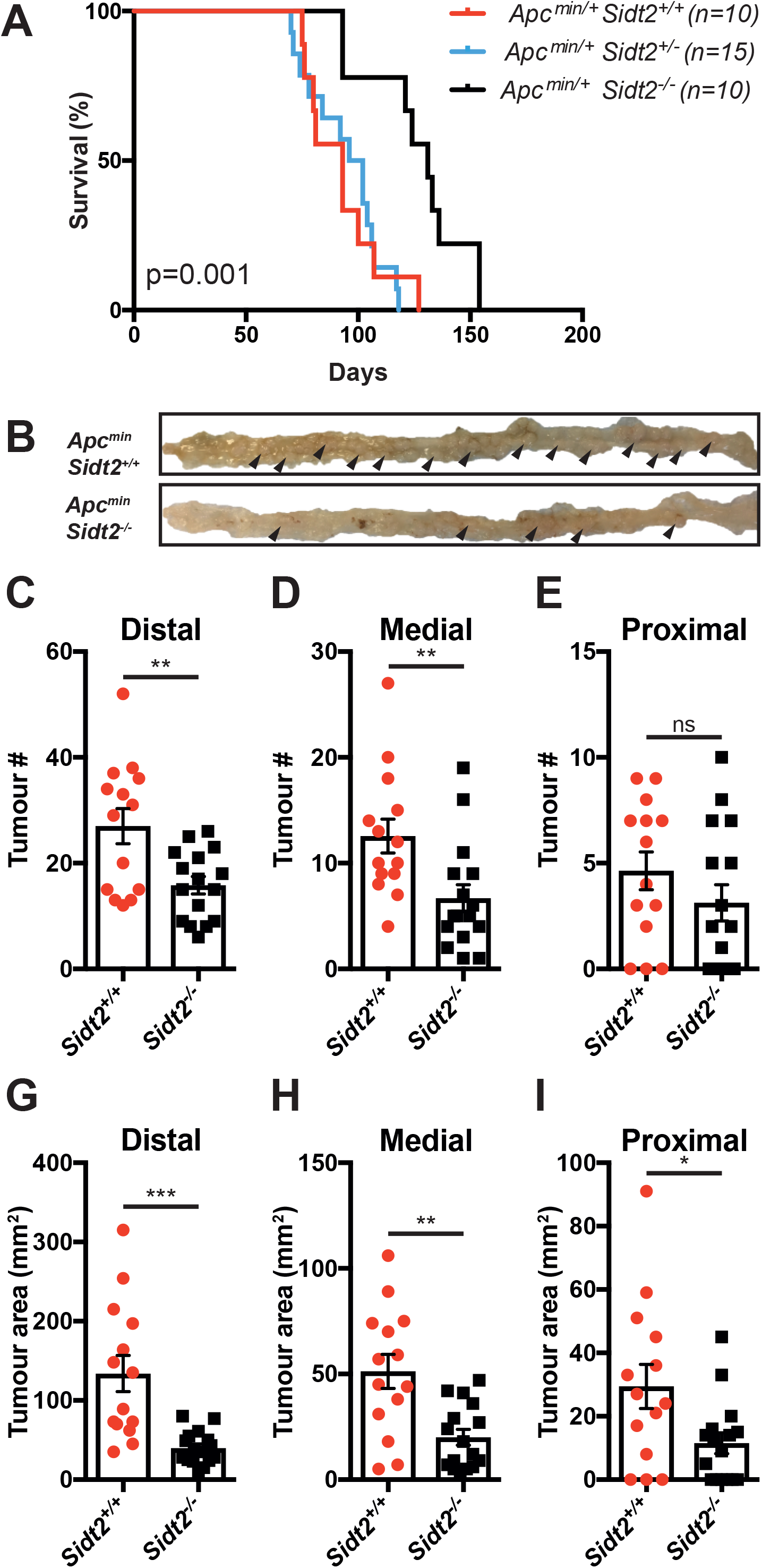
*Apc*^*min*/+^;*Sidt2*^−/−^ mice have enhanced survival and smaller tumors compared to controls. **A)** Kaplan–Meier survival curve of *Apc*^*min*/+^;*Sidt2*^+/+^, *Apc*^*min*/+^;*Sidt2*^+/−^ and *Apc*^*min*/+^;*Sidt2*^−/−^ mice. Median survival rates of *Apc*^*min*/+^;*Sidt2*^+/+^, *Apc*^*min*/+^;*Sidt2*^+/−^ and *Apc*^*min*/+^;*Sidt2*^−/−^ mice are 93, 99 and 131 days respectively. **(B)** Representative image of distal small intestine of 100-day old *Apc*^*min*/+^;*Sidt2*^+/+^; and *Apc*^*min*/+^;*Sidt2*^−/−^ mouse Total number **(C-E)** and area **(F-I)** of visible tumors in the colon and small intestine (distal, medial and proximal) was quantified in 100-day old *Apc*^*min*/+^;*Sidt2*^+/+^ (n=12) and *Apc*^*min*/+^;*Sidt2*^−/−^ (n=12) mice. Error bars represent ± SEM. * indicates *P* < 0.05, ** indicates *P* < 0.01, *** indicates *P* < 0.001 as calculated by unpaired student’s t-test.

Since *Apc*^*min*/+^;*Sidt2*^−/−^ mice had significantly smaller tumors than *Apc*^*min*/+^;*Sidt2*^+/+^ mice across all segments of the small intestine (Figure 2G-I), we hypothesized that loss of SIDT2 does not affect tumor initiation, but instead plays a role in the growth of tumors by impairing cell proliferation. To investigate this possibility, we compared expression of Ki67 in tumors of *Apc*^*min*/+^;*Sidt2*^−/−^ and *Apc*^*min*/+^;*Sidt2*^+/+^ mice (Figure 3A). Consistent with the decrease in Ki67-positive cells observed in *Kras*^*LSL-G12D*/+^;*Sidt2*^−/−^ mice, *Apc*^*min*/+^;*Sidt2*^−/−^ mice had significantly less Ki67 positive staining overall (Figure 3B), as well as fewer intratumoral Ki67-positive cells (Figure 3C) compared to *Apc*^*min*/+^;*Sidt2*^+/+^ mice.

**Figure 3.**
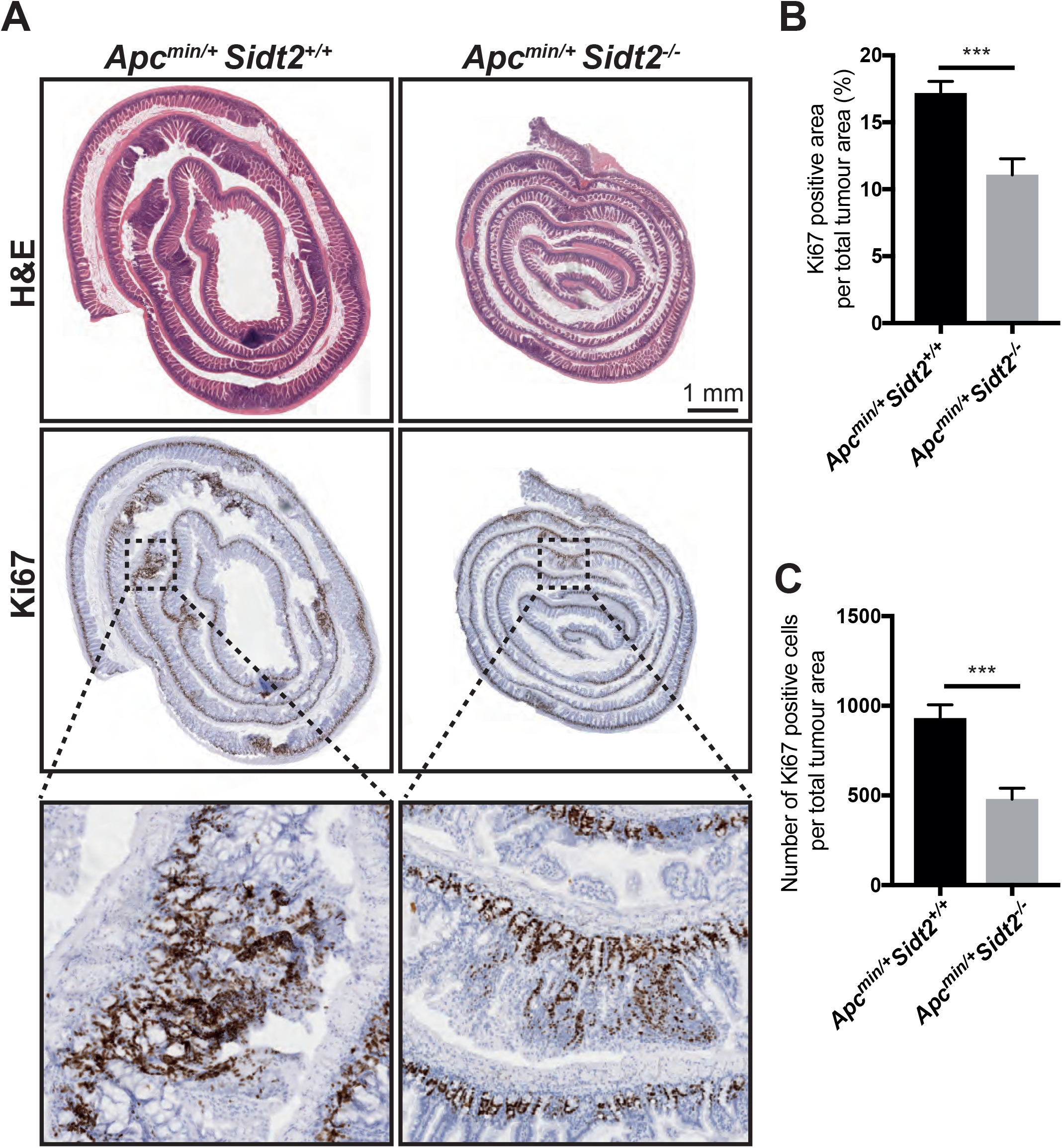
Loss of SIDT2 impairs tumor proliferation. **(A)** Representative H&E and Ki67 stained histological sections of distal small intestine *Apc*^*min*/+^;*Sidt2*^+/+^ and *Apc*^*min*/+^;*Sidt2*^−/−^ mice at 100 days of age. Quantification of **(B)** Ki67 positive staining area as a percentage of total tumor area and **(C)** average number of Ki67 positive cells per tumor. Analysis was performed on >20 high-power fields across three mice. Error bars represent ± SEM. *** indicates *P* <0.001 as calculated by unpaired student’s t-test.

### SIDT2 is required to prevent dsRNA accumulation and PKR/eIF2α pathway activation

Given the role of SIDT2 in transporting dsRNA across the endo-lysosomal membrane ^4^, we next assessed the effect of SIDT2 on the subcellular localisation of dsRNA within the DSI. To do so, we performed immunofluorescence staining on frozen sections of the DSI of 100-day old *Apc*^*min*/+^;*Sidt2*^−/−^ and *Apc*^*min*/+^;*Sidt2*^+/+^ mice using the J2 monoclonal antibody (Figure 4A), which specifically detects dsRNA helices at least 40bp in length, in a sequence-independent manner ^17^. Notably, staining for dsRNA was readily observed within the intestinal crypts of *Apc*^*min*/+^;*Sidt2*^−/−^ mice but was absent in crypts of *Apc*^*min*/+^;*Sidt2*^+/+^ animals and in other parts of the DSI, including tumors (Figure 4B). To confirm that this was not specific to the *Apc*^*min*/+^ mouse model, we also performed dsRNA staining in lungs of *Kras*^*LSL-G12D*/+^; *Sidt2*^+/+^ and *Kras*^*LSL-G12D*/+^; *Sidt2*^−/−^ mice using immunohistochemistry (Figure S3A). Consistent with our previous data, we observed a significant increase in cytosolic dsRNA staining in *Kras*^*LSL-G12D*/+^; *Sidt2*^−/−^ mice compared to controls (Figure S3B) suggesting that loss of SIDT2 leads to accumulation of dsRNA in the cytosol.

**Figure 4.**
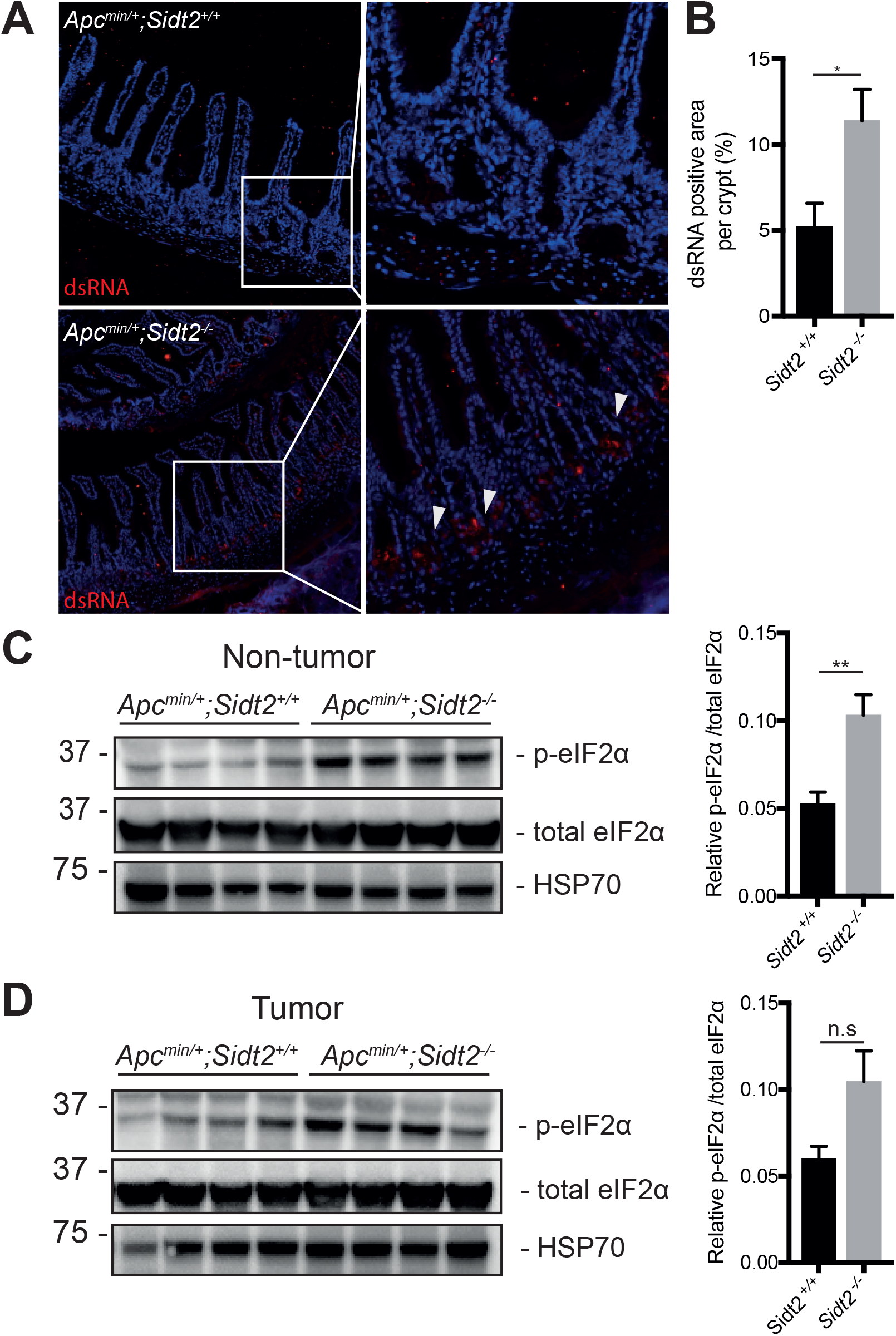
Loss of SIDT2 leads to dsRNA accumulation within intestinal crypts and increased phosphorylation of eIF2α. **(A)** Representative image of dsRNA staining of distal small intestine of 100-day old *Sidt2*^+/+^;*Apc*^*min*/+^ and *Sidt2*^−/−^;*Apc*^*min*/+^ mice. **(B)** Quantification of dsRNA-positive stained area per crypt (n = 3-4 mice per genotype). Western blot and densitometry analysis of eIF2α phosphorylation in non-tumor **(C)** and tumor **(D)** tissue from distal small intestine of 100-day old *Sidt2*^+/+^;*Apc*^*min*/+^ and *Sidt2*^−/−^;*Apc*^*min*/+^ mice normalized to total eIF2α. Each lane corresponds to an individual mouse (n = 4 mice per genotype). Error bars represent ± SEM. * indicates *P* < 0.05, ** indicates *P* < 0.01 as calculated by unpaired student’s t-test.

Cytosolic dsRNAs are bound by RNA-dependent protein kinase (PKR), leading to its auto-phosphorylation and activation ^18^. Activated PKR subsequently phosphorylates the α subunit of protein synthesis initiation factor eIF2 (eIF2α), resulting in inhibition of protein translation as well as anti-viral and anti-tumor effects (10, 11). We therefore wished to determine whether the accumulation of dsRNA in cells deficient for SIDT2 was likely to activate PKR. Although we tested multiple antibodies raised against phosphorylated PKR, none showed an ability to recognize activated murine PKR in control samples via western blot (data not shown), so instead we compared phosphorylation of eIF2α as a proxy for PKR activation. Moreover, since we only observed SIDT2-dependent dsRNA accumulation in the intestinal crypts and not in tumors themselves, we analyzed p-eIF2α expression separately in non-tumor and tumor tissue from the DSI of *Apc*^*min*/+^;*Sidt2*^−/−^ and *Apc*^*min*/+^;*Sidt2*^+/+^ mice. Notably, *Apc*^*min*/+^;*Sidt2*^+/+^ mice displayed higher p-eIF2α levels in non-tumor tissue (Figure 4C), consistent with dsRNA accumulation and activation of PKR within these cells. This SIDT2-dependent increase in p-eIF2α was less apparent in tumor tissue (Figure 4D).

In addition to its role in the inhibition of protein translation, PKR activation has been shown to mediate cellular stress responses via regulation of mitogen-activated protein kinases (MAPK) such as c-Jun n-terminal kinase (JNK) ^19–21^ and promote apoptosis via caspase 8 and NFκB ^22^. In line with our hypothesis that PKR activation is increased in the intestinal crypts in the absence of SIDT2, we observed a concurrent increase in phosphorylation of JNK in normal intestinal tissue of *Apc*^*min*/+^;*Sidt2*^−/−^ mice compared to *Apc*^*min*/+^;*Sidt2*^+/+^ mice (Figure 5A-B). To assess whether loss of SIDT2 leads to increased caspase 8-mediated apoptosis, we assessed and compared cleavage of caspase 8 in the DSI of *Apc*^*min*/+^;*Sidt2*^−/−^ and *Apc*^*min*/+^;*Sidt2*^+/+^ mice, and observed an increase in caspase 8 cleavage products in *Apc*^*min*/+^;*Sidt2*^−/−^ normal intestinal tissue (Figure 5A, C). We next performed immunohistochemical staining on intestinal swiss rolls of *Apc*^*min*/+^;*Sidt2*^−/−^ and *Apc*^*min*/+^;*Sidt2*^+/+^ mice (Figure 5D), which revealed an increased number of cleaved-caspase 3 positive cells within intestinal crypts lacking SIDT2 (Figure 5E). This was further confirmed via TUNEL staining in which *Apc*^*min*/+^;*Sidt2*^−/−^ showed increased TUNEL-positive cells within intestinal crypts compared to controls (Figure 5F). Taken together, these data strongly imply that loss of SIDT2 leads to increased caspase 8 and 3-mediated apoptosis within intestinal crypts, consistent with the restricted tumor growth observed in *Apc*^*min*/+^;*Sidt2*^−/−^ animals.

**Figure 5.**
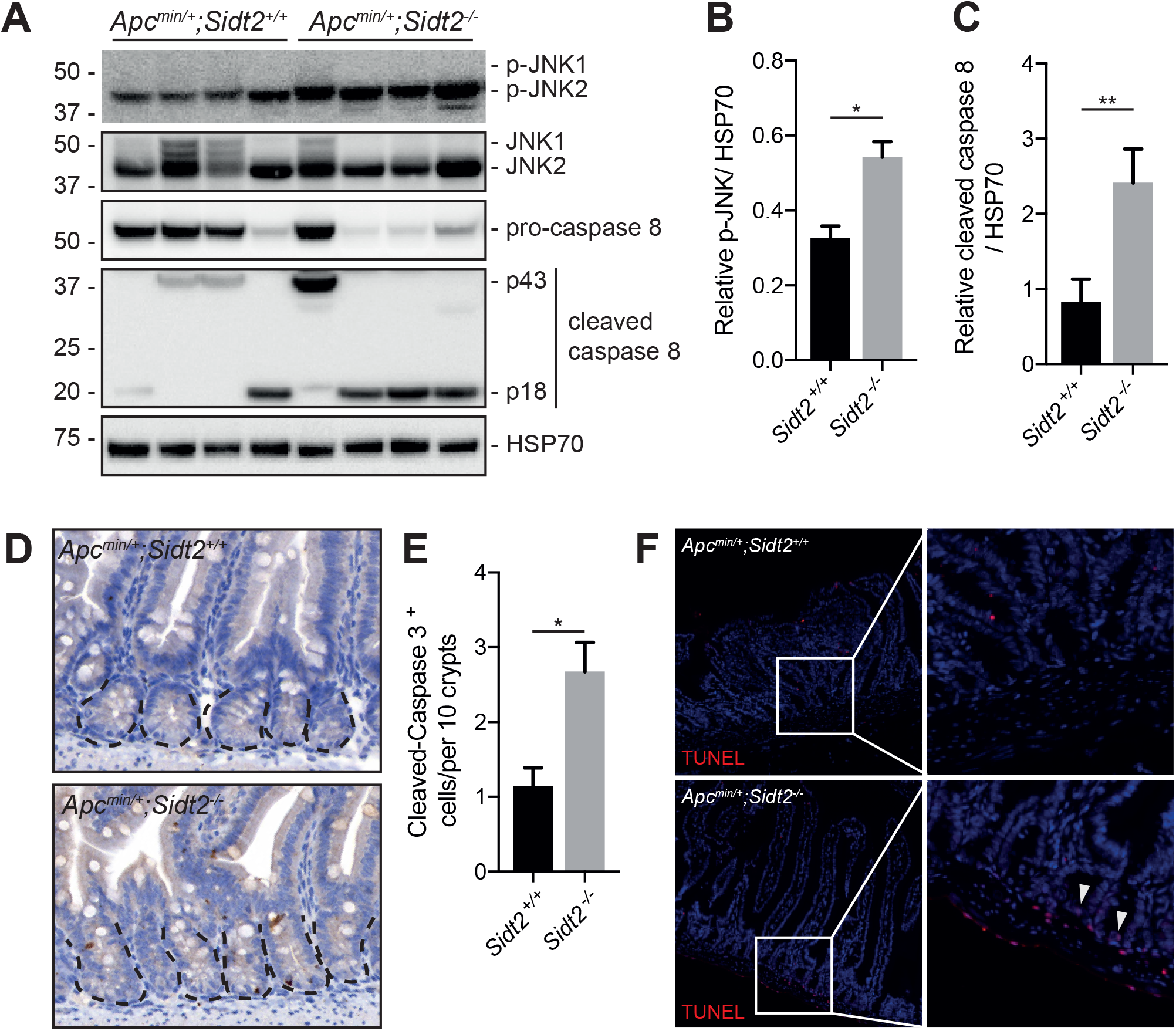
*Sidt2*^−/−^;*Apc*^*min*/+^ mice have increased apoptosis in intestinal tissue. Western blot analysis **(A)** Phosphorylation of JNK and cleavage of caspase 8 was assessed in normal intestinal tissue of 100-day old *Sidt2*^+/+^;*Apc*^*min*/+^ and *Sidt2*^−/−^;*Apc*^*min*/+^ mice. Each lane corresponds to an individual mouse (n = 4 mice per genotype). Densitometry quantification of **(B)** p-JNK and **(C)** cleaved caspase 8 normalised to HSP70. Error bars represent ± SEM. * indicates *P* < 0.05, ** indicates *P* < 0.01 as calculated by unpaired student’s t-test. **(D)** Representative images of cleaved-caspase 3 staining of distal small intestine of 100-day old *Sidt2*^+/+^;*Apc*^*min*/+^ and *Sidt2*^−/−^;*Apc*^*min*/+^ mice. **(E)** Quantification of dsRNA-positive stained area per crypt (n = 7 mice per genotype). Error bars represent ± SEM. * indicates *P* < 0.05, as calculated by unpaired student’s t-test. **(F)** Representative images of TUNEL staining of positive cells from distal small intestine of 100-day old *Sidt2*^+/+^;*Apc*^*min*/+^ and *Sidt2*^−/−^;*Apc*^*min*/+^ mice, n = 3 animals per genotype.

### Lower *SIDT2* expression is associated with improved survival in different human cancers

Finally, to explore the role of SIDT2 in human cancer, we determined whether different levels of intratumoral *SIDT2* expression are associated with changes in patient survival. Using data collected by The Cancer Genome Atlas Research Network (http://cancergenome.nih.gov/) and analyzed via The Pathology Atlas ^23^, we observed that lower intratumoral *SIDT2* RNA levels were associated with significantly improved prognosis in 5 of 17 different cancers (renal, thyroid, gastric, glioma, urothelial) (Figure 6A-E). However, consistent with a context-dependent role, intratumoral *SIDT2* RNA levels showed no prognostic significance in 10 other cancers (including LUAD and colon cancer), and lower intratumoral *SIDT2* levels were actually associated with poorer survival in pancreatic and cervical cancers (Figure S5).

**Figure 6.**
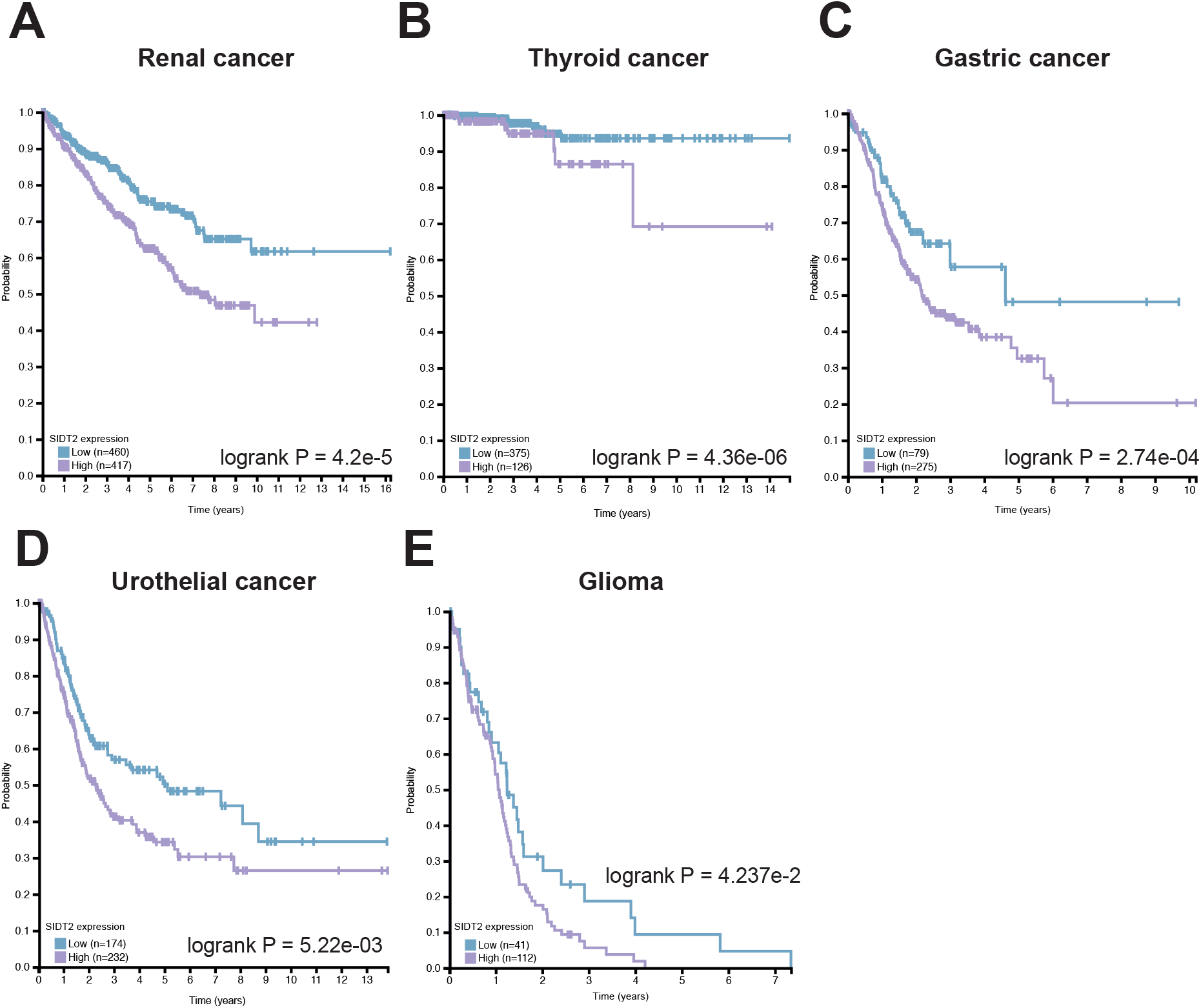
Lower SIDT2 expression is associated with improved survival in some cancers. Kaplan–Meier curves of overall survival of **(A)** renal, **(B)** thyroid, **(C)** gastric, **(D)** urothelial cancer and **(E)** glioma patients stratified against *SIDT2* expression from publicly available RNAseq data. The results shown are in whole based upon data generated by the TCGA Research Network: http://cancergenome.nih.gov/.

## DISCUSSION

In this study, we show that the mammalian SID-1 ortholog SIDT2, which is a transcriptional target for p53, promotes tumor growth. Specifically, loss of SIDT2 in mice caused reduced tumor burden and enhanced survival, with the latter observation supported by data from patients with several different cancers in which lower *SIDT2* expression was associated with an improved prognosis.

Mechanistically, our data from *Apc*^*min*^ animals suggest a model in which the absence of SIDT2 leads to impaired RNautophagy, accumulation of intracellular dsRNA, increased phosphorylation of eIF2α and JNK, and finally increased apoptosis via activation of caspase 8 (Figure 7). Notably, when visualized, these changes were only observed in the intestinal crypts and not in tumors themselves, and is potentially in keeping with the strong expression of SIDT2 observed within the crypts of human small intestine ^23^ (Figure S4). Nevertheless, such observations invite the question of how such crypt-related changes could impact subsequent tumor growth. One possible explanation is that the increased apoptosis observed in SIDT2-deficient mice affects cancer stem cells, which reside within the intestinal crypt and play a key role in subsequent tumor growth. Consistent with this, selective depletion of intestinal stem cells has previously been shown to restrict primary tumor growth in mice ^24^ and, this would be in keeping with our observation that SIDT2-deficient tumors show reduced proliferation (Figure 3). Another possible explanation is that increased eIF2α activation in the absence of SIDT2 induces differentiation (and thus loss) of these same intestinal stem cells, as has been observed by others ^25^. Further work to investigate the specific effects of SIDT2 on cancer stem cell development will hopefully shed light on these possibilities.

**Figure 7.**
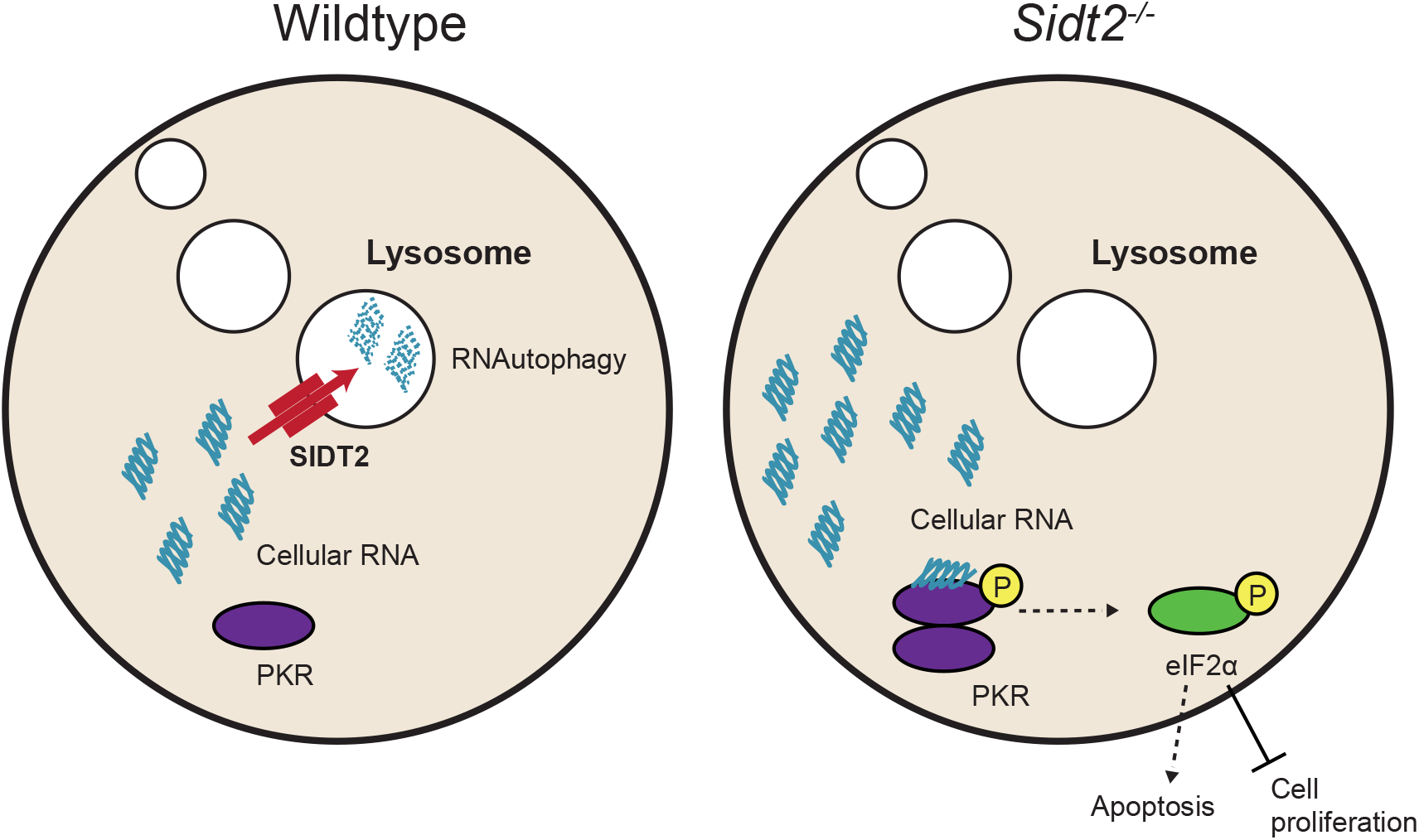
Loss of SIDT2-mediated RNAutophagy leads to increased apoptosis and cell proliferation. SIDT2 is a dsRNA transporter mediates the lysosomal degradation of cellular RNAs via RNAutophagy. Loss of SIDT2 leads to accumulation of cellular RNA within the cytosol. Binding of these RNAs by protein kinase RNA-activated (PKR), leading to its activation and subsequent phosphorylation of eukaryotic translation initiation factor 2 alpha (eIF2α). Phosphorylation of eIF2α leads to inhibition of protein translation and cell proliferation, as well as induction of apoptosis.

Although we were unable to directly test PKR activation due to a lack of suitable reagents, PKR nonetheless seems a likely candidate in mediating the downstream effects in our *Apc*^*min*^ *Sidt2*^−/−^ mice. After all, PKR has well-established roles not only in binding and responding to intracellular dsRNA, but also inducing eIF2α phosphorylation, JNK activation and caspase 8-mediated activation of caspase 3 to induce apoptosis. Consistent with our results, PKR expression and auto-phosphorylation has previously been reported to be increased in multiple human malignancies, including colon and lung cancer ^26,27^, and those with higher p-PKR and p-eIF2α had significantly longer survival ^28–30^.

Given the presence of intracellular dsRNA in *Apc*^*min*^;*Sidt2*^−/−^ intestinal crypts, we also assessed whether there was induction of a type I interferon (IFN) response in these tissues. However, we were unable to detect upregulation of *Ifnβ* or various IFN stimulated genes in tissues of *Apc*^*min*/+^;*Sidt2*^−/−^ mice (data not shown), suggesting that the reduced tumor burden observed in SIDT2-deficient mice is unlikely to be due to the anti-tumor effects of type I IFN ^31^. This lack of a type I IFN response may also provide a clue as to the nature of the dsRNA that accumulates in the absence of SIDT2. Specifically, the type I IFN response to cytoplasmic dsRNA is mainly orchestrated by the RIG-I-like receptors (RLRs), RIG-I and MDA-5 ^32,33^. RIG-I specifically recognizes short dsRNA and ssRNA with 5’ triphosphate ends ^34,35^ - a common feature of viral RNAs - and MDA-5 is critical for the detection of long dsRNAs (>1000 bp) ^36–38^. The lack of a detectable type I IFN response in *Apc*^*min*/+^;*Sidt2*^−/−^ mice therefore suggests that the dsRNAs which accumulate in the intestinal crypts of these animals do not possess 5’triphopshate ends and are not long enough to activate MDA-5. In contrast, PKR requires a minimum dsRNA region of only 30 bp ^39,40^ and can recognize RNAs with limited stem loop or duplex structures ^41^, including siRNAs ^42,43^, snoRNAs ^44^ and bacterial RNAs ^45^. Indeed, PKR has recently been reported to bind endogenous nuclear dsRNAs during cell mitosis ^21^, noncoding Alu RNA and mitochondrial RNAs that are capable of forming intramolecular dsRNA structures ^46^. These RNA species may therefore be more likely to accumulate in the absence of SIDT2. At the same time, it has been proposed that autophagy is important for the degradation of many types of cytosolic RNAs, including retrotransposon, viral and cellular messenger RNAs ^6,47,48^ so impaired lysosomal degradation of RNA in the absence of SIDT2 could also lead to accumulation of these RNAs within the intestinal crypt. Future studies to identify the RNA cargo of SIDT2 within the intestine will clarify these possibilities.

Finally, it should be noted that the apparent protective effect of lower intratumoral *SIDT2* levels on patient survival was limited to certain types of cancers. Moreover, in pancreatic and cervical cancers, lower intratumoral *SIDT2* levels were associated with poorer survival. Thus, the role of SIDT2 in promoting tumor growth may only apply in specific contexts, and this may explain the apparent contradiction between the data described here and our earlier findings, which found that SIDT2 functions as a tumor suppressor in a fibrosarcoma model ^12^. Regardless, the results from this study identify a role for SIDT2 and RNautophagy in promoting cancer development *in vivo*, and suggest the possibility that strategies to inhibit SIDT2 and/or RNautophagy could be a useful adjunct to existing treatment for certain types of cancer.

## AUTHOR CONTRIBUTIONS

K.C.P., T.A.N., K.T.B, T.L.P, I.P.W, L.D.A. designed experiments and/or analyzed the data. T.A.N. and K.T.B. performed experiments. K.C.P. and T.A.N. wrote the manuscript. K.C.P., L.D.A. and I.P.W. supervised the project.

## EXPERIMENTAL PROCEDURES

### Mice

Mice were bred and maintained in the animal facilities at the Walter and Eliza Hall Institute of Medical Research (WEHI) and at the Department of Radiation Oncology, Stanford University, according to national and institutional guidelines for animal care. Mice were housed in individual ventilated cages at 19 - 24 °C and maintained on a 14 h light to 10 h dark cycle with continuous access to Barastoc Custom Mixed Ration (irradiated) and acidified and filtered water. All experimental procedures were approved by the relevant animal ethics committees at the WEHI and Stanford University.

### Analysis of *Kras*^*LSL-G12D*/+^ LUAD mouse model

*Kras*^*LSL-G12D*/+^;*Sidt2*^−/−^ and *Kras*^*LSL-G12D*/+^;*Sidt2*^+/+^ aged 6-8 weeks were inoculated with 4×10^6^ PFU Adenovirus containing Cre recombinase (University of Iowa Viral Vector Core) by intra-tracheal intubation ^14^. Eighteen weeks after inoculation, the lungs were harvested, fixed in 10% formalin, and bread-loafed into approximately 12 pieces per lung. The pieces were sectioned, stained with hematoxylin and eosin (H&E) as described below, and analyzed for tumor number and tumor burden.

### Histological analyses

For histological examination of the intestine, colon, distal, medial and proximal portions of the small intestine were slit open longitudinally and the intraluminal contents removed. Each segment was then rolled up from the distal end to the proximal end using a “Swiss-rolling” technique ^49^. Preparations were then fixed in 10% (w/v) neutral-buffered formalin for at least 12 hours before embedding in paraffin, sectioned (3 µm), and prepared for general staining and immunohistochemistry (IHC). For frozen sections, Swiss roll preparations were frozen and embedded in Optimal Cutting Temperature (OCT) using a PrestoCHILL system (Milestone), and sectioned (10 µm) for immunofluorescence staining. For H&E staining, slides were incubated with hematoxylin to stain nuclei (5 min), washed in dH_2_O (1 min), incubated in 0.3% (v/v) acid ethanol to de-stain (1 dip), rinsed in dH_2_O followed by Scott's tap water, then incubated in eosin (1 min). Slides were then rinsed in dH_2_0, dehydrated, cleared and mounted.

### Immunohistochemistry and TUNEL staining

Automated staining was performed using the Dako Omnis EnVision G2 template. Dewax was performed with Clearify Clearing Agent (Dako) for 15 minutes, and antigen retrieval was done with EnVision FLEX TRS, High pH (Dako) at 97°C for 30 minutes. Antibody against Ki67 (Cell Signaling #9129) and cleaved Caspase-3 (Asp175) (R&D Systems) was diluted 1:100 in EnVision Flex Antibody Diluent (Dako) and incubated at 32°C 60 minutes. Envision+ System-HRP Labelled Polymer Anti-Rabbit antibodies (Dako) were incubated for 20 min at 32°C. Slides were counter-stained with Mayer Hematoxylin, dehydrated, cleared, and mounted with MM24 Mounting Medium (Surgipath-Leica). Tissue sections were imaged using the 3D Histech Slide Scanner and quantification analysis of Ki67 and cleaved-caspase 3positive cells was performed using ImageJ. For TUNEL staining, slides were prepared and stained using ApopTag® Red In Situ Apoptosis Detection Kit (Merk Millipore) according to manufacturer’s instructions and imaged on a Zeiss Axio Observer wide-field fluorescence microscope.

### Immunofluorescence staining of dsRNA

Frozen sections were fixed in 4% paraformaldehyde for 10 min on ice and washed 3 times in cold PBS permeabilized using 0.1% Tween/PBS, blocked in 5% normal goat serum for 1 h and stained with J2 (SciCon) primary antibody (1:200) overnight. Slides were subsequently washed 3 times for 5 min in PBS and then incubated with anti-mouse Alexa-594 secondary antibody for 1 h at room temperature. Sections were washed 3 times for 5 min in PBS and then cover-slipped and mounted using Fluoroshield mounting medium with DAPI (Sigma Aldrich). Stained sections were imaged on a Zeiss Axio Observer wide-field fluorescence microscope.

### Western blotting

Intestinal tumor and adjacent non-tumor tissue were lysed in KALB lysis buffer supplemented with protease inhibitor and phosphatase inhibitor (Roche) and then homogenized using a QIAGEN TissueLyser II, according to manufacturer’s instructions. Homogenates were subsequently cleared by centrifugation at 13,000 rpm at 4°C for 20 minutes and denatured in 4x SDS PAGE sample buffer at 95°C for 5 min. 60µg of protein lysate was separated on NUPAGE Novex 4-12% Bis-Tris Gels (Life Technologies), and transferred electrophoretically to PVDF membranes, blocked in 5% BSA/TBST, incubated with the relevant primary antibodies overnight. Membranes were incubated for 1 h with horseradish peroxidase-conjugated secondary antibodies, washed, treated with Luminata Forte Western HRP substrate (Millipore), and visualized on the ChemiDoc MP system (Bio-Rad). Prmary antibodies used were: total eiF2α (Cell Signaling, #9722, 1:1000), phospho-eIF2α (Cell Signaling, # 9721, 1:1000), phospho-JNK (Cell Signaling, #4668P, 1:1000), total JNK (Cell Signaling, #9252, 1:1000), pro-caspase 8 (WEHI in-house antibody, 1:1000), mouse specific cleaved caspase-8 (Asp387) (Cell Signaling #9429, 1:1000), Hsp70 (Thermo Fisher Scientific, #MA3-007, 1:10 000).

### Analysis of *SIDT2* expression on cancer patient survival

Maximally separated Kaplan–Meier curves of overall survival of cancer patients stratified against *SIDT2* expression were generated from data by the TCGA Research Network: http://cancergenome.nih.gov/ and accessed via The Human Protein Atlas ^23^. Based on the FPKM value of SIDT2, patients were classified into two expression groups and the correlation between expression level and patient survival was examined.

### Statistical analysis

Statistical analyses were performed in GraphPad Prism 7 software using unpaired, two-tailed student’s t-tests, except in the case of the zsurvival analysis, where a generalized Wilcoxon (Gehan–Breslow) test was used to compare survival curves. *P* values < 0.05 were considered statistically significant.

