## Supplementary figures and images for "SIDT2 RNA transporter promotes lung and gastrointestinal tumor development"

### Figure S1

#### Supplemental Figure 1

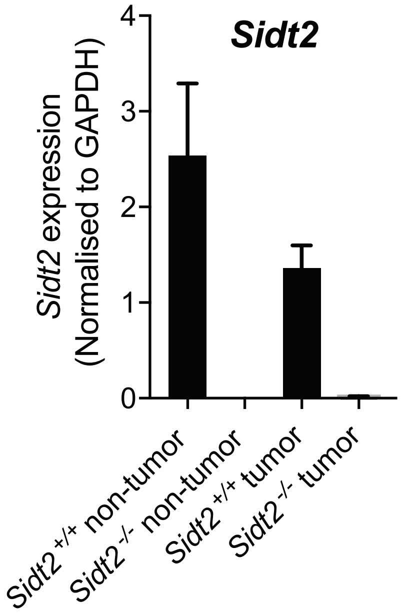

### Figure S2

#### Supplemental Figure 2

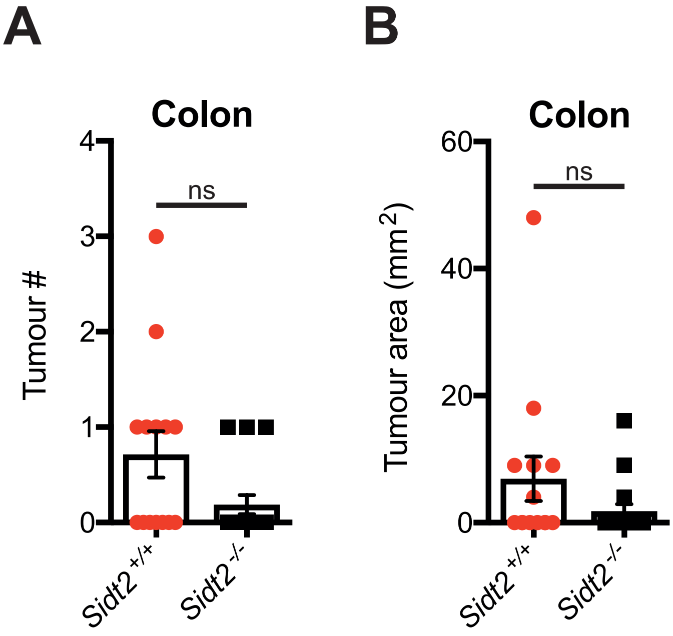

### Figure S3

#### Supplemental Figure 3

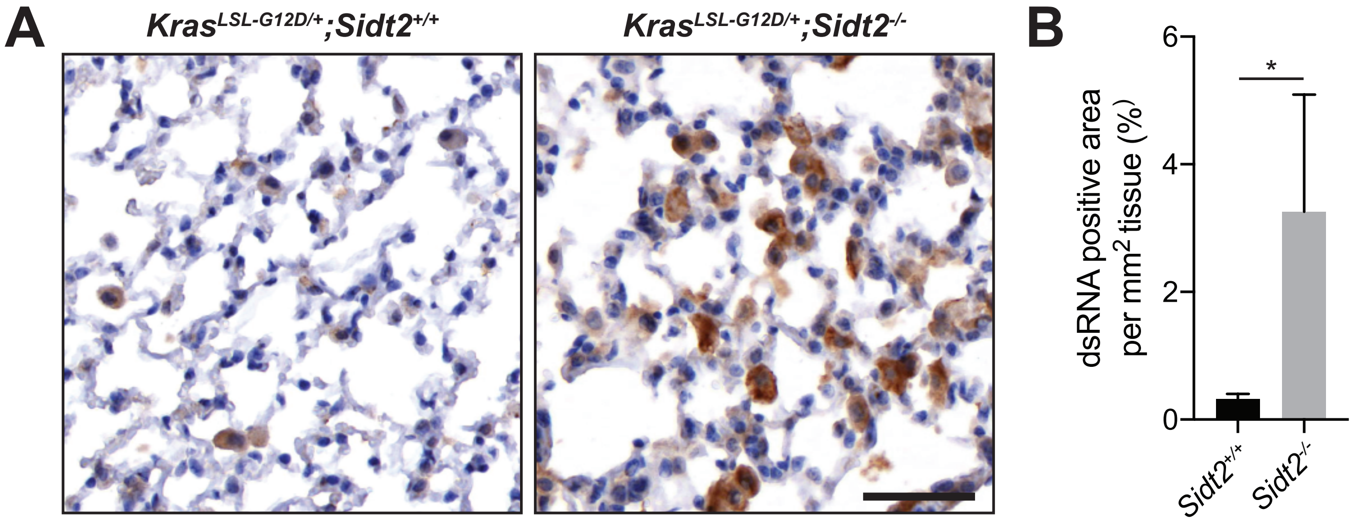

### Figure S4

#### Supplemental Figure 4

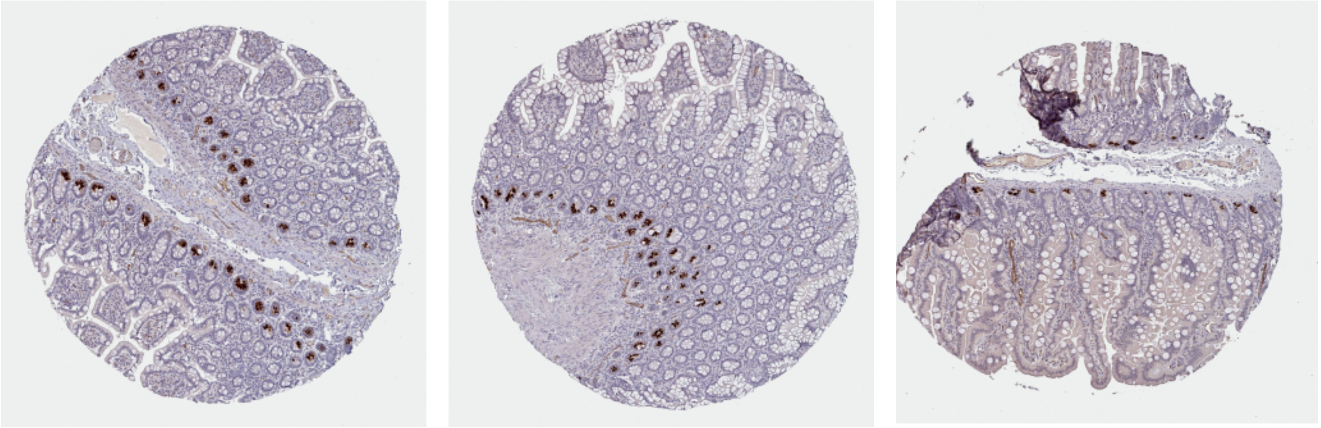

### Figure S5

#### Supplemental Figure 5

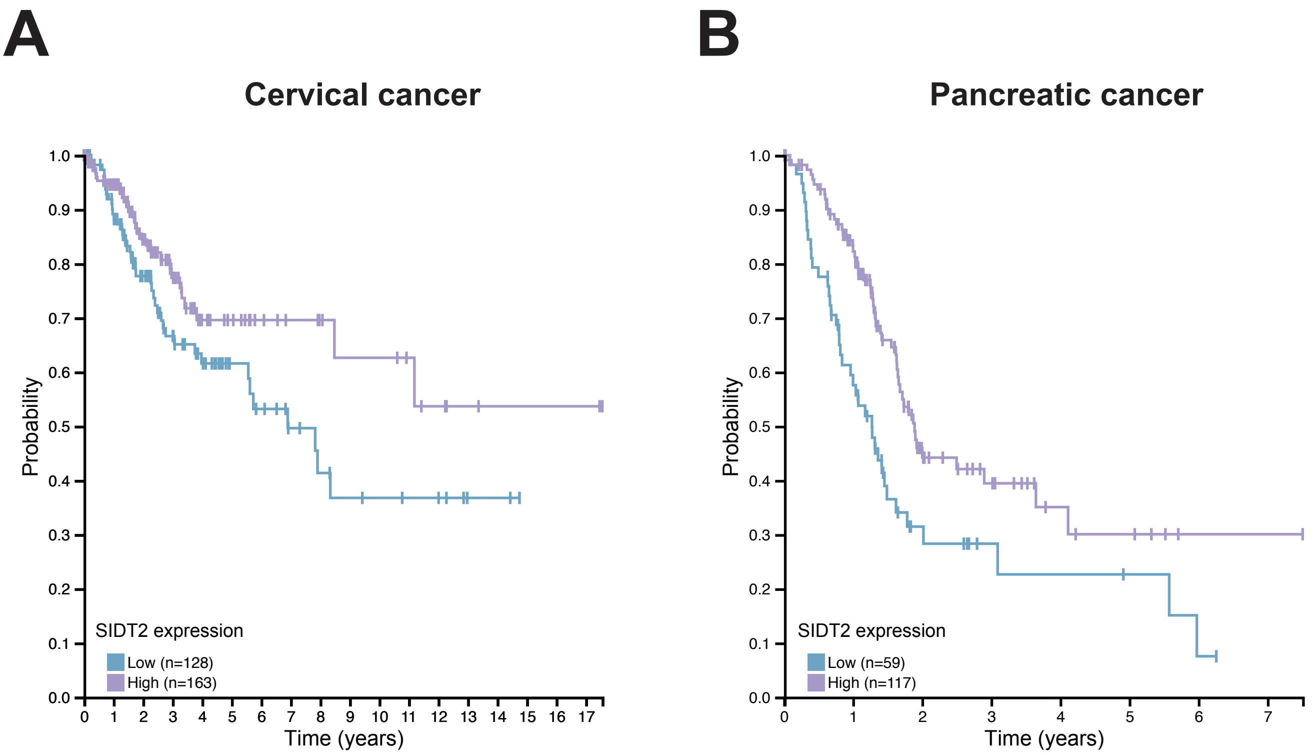
