## Supplemental Figure Legends for "SIDT2 RNA transporter promotes lung and gastrointestinal tumor development"

**Supplemental Figure 1 – *Sidt2* is expressed in normal intestinal and tumour tissue in the distal small intestine of *Apc^min/+^* mice.**

*Sidt2* expression in tumour and adjacent non-tumour tissue isolated from 100-day old *Sidt2^+/+^;Apc^min/+^* and *Sidt2^-/-^;Apc^min/+^* mice assessed via qRT-PCR. n = 5 mice per group. Data is plotted as mean ± SEM.

**Supplemental Figure 2 – Loss of SIDT2 does not affect tumor burden in *Apc^min/+^* mouse model**

Total number **(A)** and area **(B)** of visible tumors in the colon and small intestine (distal, medial and proximal) was quantified in 100-day old *Apc^min/+^;Sidt2^+/+^* (n=12) and *Apc^min/+^;Sidt2^-/-^* (n=12) mice. Error bars represent ± SEM.

**Supplemental Figure 3 – Loss of SIDT2 leads to accumulation of cytosolic dsRNA in the lungs of *Kras^LSL-G12D/+^* ; *Sidt2^-/-^* mice.**

**(A)** Representative image of dsRNA staining of lung sections from *Kras^LSL-G12D/+^* ; *Sidt2^+/+^* and *Kras^LSL-G12D/+^* ; *Sidt2^-/-^* mice. **(B)** Quantification of dsRNA-positive stained area per non-tumour area (n = 6 mice per genotype). Error bars represent ± SEM. * indicates *P* < 0.05 as calculated by Mann Whitney non-parametric test.

**Supplemental Figure 4 – Human SIDT2 is expressed in glandular crypts of small intestine**

Publicly accessible images from The Human Protein Atlas of immunohistochemical stained sections of human small intestine from 3 individual healthy patient samples shows strong cytoplasmic localisation of SIDT2 in glandular cells (Uhlen et al., 2017).

**Supplemental Figure 5 – Lower SIDT2 expression is associated with lower survival for pancreatic and cervical cancer**

Kaplan–Meier curves of overall survival of **(A)** cervical and **(B)** pancreatic cancer patients stratified against *SIDT2* expression from publicly available RNAseq data. The results shown are based upon data generated by the TCGA Research Network: http://cancergenome.nih.gov/.
